# Host factors drive the within-host competition between highly and low pathogenic H5N8 avian influenza viruses

**DOI:** 10.1101/2021.04.06.438580

**Authors:** Pierre Bessière, Thomas Figueroa, Amelia Coggon, Charlotte Foret-Lucas, Alexandre Houffschmitt, Maxime Fusade-Boyer, Gabriel Dupré, Jean-Luc Guérin, Maxence Delverdier, Romain Volmer

## Abstract

Highly pathogenic avian influenza viruses (HPAIV) emerge from low pathogenic avian influenza viruses (LPAIV) through the introduction of basic amino acids at the hemagglutinin (HA) cleavage site. Following viral evolution, the newly formed HPAIV likely represents a minority variant within the index host, predominantly infected with the LPAIV precursor. Using reverse-genetics engineered H5N8 viruses differing solely at the HA cleavage, we tested the hypothesis that the interaction between the minority HPAIV and the majority LPAIV could modulate the risk of HPAIV emergence and that the nature of the interaction could depend on the host species. In chickens, we observed that the H5N8_LP_ increased H5N8_HP_ replication and pathogenesis. By contrast, the H5N8_LP_ antagonized H5N8_HP_ replication and pathogenesis in ducks. Ducks mounted a more potent antiviral innate immune response than chickens against the H5N8_LP_, which correlated with H5N8_HP_ inhibition. These data provide experimental evidence that HPAIV may be more likely to emerge in chickens than in ducks and underscore the importance of within-host viral variants interactions in viral evolution.

## INTRODUCTION

Highly pathogenic avian influenza virus (HPAIV) outbreaks have a major impact on animal health, food security and economy, as well as on public health (EFSA Panel on Animal Health and Welfare (AHAW) et al., 2017; Lee et al., 2021). A better understanding of the factors leading to HPAIV emergence is therefore of paramount importance. Avian influenza viruses of H5 and H7 subtypes can become highly pathogenic through the introduction of multiple basic amino acids within the hemagglutinin (HA) cleavage site (Böttcher-Friebertshäuser et al., 2014). Several mechanisms account for these introductions: nucleotide substitutions and insertions or non-homologous recombination with viral or cellular RNAs (Abdelwhab et al., 2013; Lee et al., 2021; Richard et al., 2017). The acquisition of a multi-basic cleavage site (MBCS) is a virus-dependent event that occurs in a bird infected with a parental low pathogenic avian influenza virus (LPAIV). To emerge successfully, the newly formed HPAIV must become a predominant variant in order to overcome the transmission bottleneck between individuals (Hughes et al., 2012; McCrone and Lauring, 2018; Varble et al., 2014). In order to achieve this, the newly formed HPAIV must therefore outcompete its LPAIV precursor within the individual in which it has arisen. We therefore suggest that HPAIV emergence is a two-step process: firstly, the acquisition of a MBCS, followed by the ability to become a predominant variant of the viral quasi-species within an individual.

Over the past decades, the vast majority of HPAIV emergences have been linked to H5 and H7 LPAIV circulation in *Galliformes* (such as chickens and turkeys) (Abdelwhab et al., 2013; EFSA Panel on Animal Health and Welfare (AHAW) et al., 2017; Lee et al., 2021). By contrast, *Anseriformes* (such as ducks and geese) are considered as reservoirs for LPAIV precursors, rather than a species in which HPAIV emerge (Bodewes and Kuiken, 2018; Munster et al., 2007; van Dijk et al., 2018; Wahlgren, 2011). These observations suggest that host factors could modulate HPAIV emergence. To our knowledge, this has never been investigated experimentally.

To model the intra-host competition between a newly formed HPAIV and its parental LPAIV, we co-infected embryonated eggs and chickens and ducks *in vivo* with a H5N8 HPAIV as a minority variant and a reverse-genetics engineered LPAIV that differed from the HPAIV only at the level of the HA cleavage site, as a majority variant. Our results demonstrate that chickens and ducks have opposite effects on the interaction between the H5N8 HPAIV and LPAIV and that the HPAIV has a stronger selective advantage in chickens than in ducks. To our knowledge this is the first experimental evidence that HPAIV emergence could be more likely in chickens than in ducks.

## RESULTS

### Characterization of H5N8_HP_ and H5N8_LP_ viruses in cell culture

We used reverse-genetics to generate the wild-type H5N8_HP_ from a HPAIV isolated during the 2016 H5N8 epizootics in France (Guinat et al., 2018). This virus belonged to clade 2.3.4.4 group B, which caused high levels of mortality in *Galliformes* and in wild and domestic *Anseriformes* during the 2016-2017 HPAIV outbreak in Europe (Grund et al., 2018; Guinat et al., 2018; Kleyheeg et al., 2017; More et al., 2017). Using site-directed mutagenesis and reverse-genetics, we mutated the H5N8_HP_ HA polybasic cleavage site PLRELRRLR/G to a monobasic cleavage site PQRETR/G to obtain the H5N8_LP_ virus with a typical LPAIV HA sequence (NCBI Genbank accession number: AB261853.1) (Hiono et al., 2016). The H5N8_HP_ and the H5N8_LP_ differed solely at the level of the HA cleavage site, as verified by whole genome sequencing following virus amplification in chicken embryonated eggs (Fig. 1A).

**Fig. 1.**
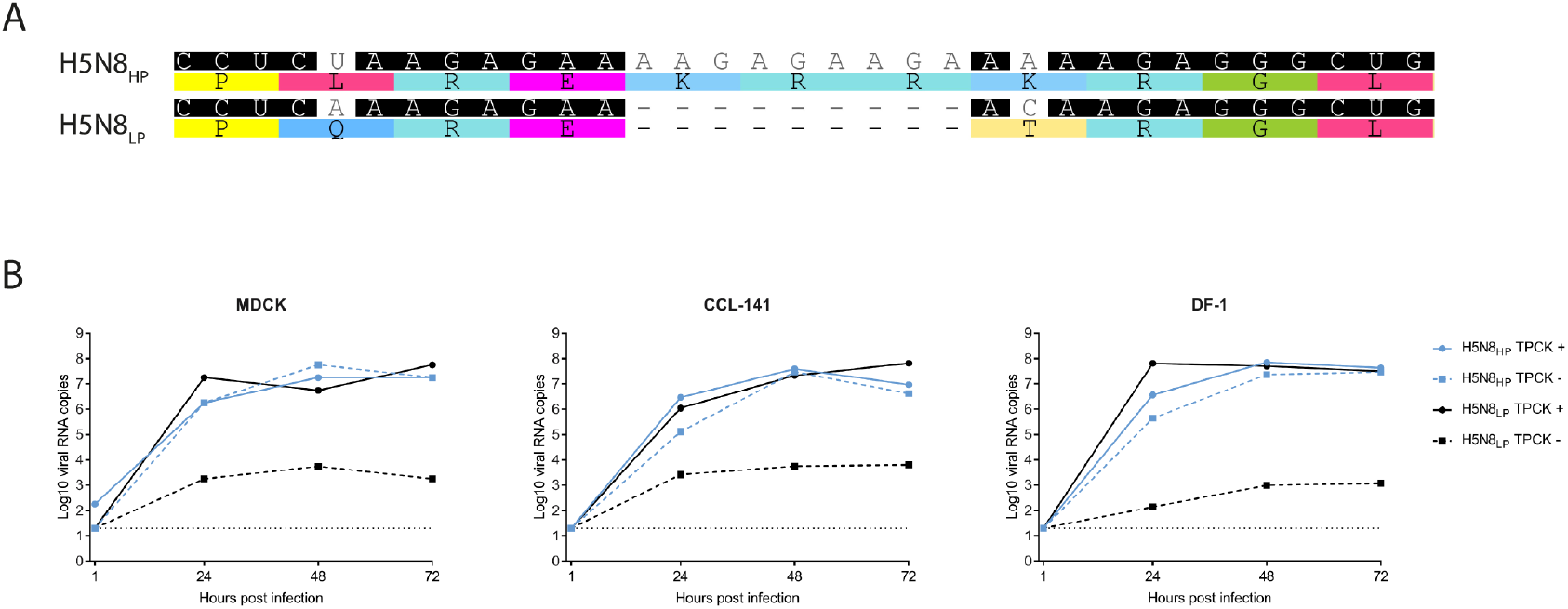
Characterization of H5N8_HP_ and H5N8_LP_ viruses in cell culture. **(A)** Sequence alignment of H5N8_HP_ and H5N8_LP_ HA cleavage sites. **(B)** H5N8_HP_ and H5N8_LP_ growth kinetics in the absence or presence of exogenous trypsin in Madin-Darby canine kidney (MDCK) cells, DF-1 cells and CCL-141 cells. Cells were infected at a MOI of 10^−5^ TCID_50_. HA RNA load was analyzed by RT-qPCR using primers common to H5N8_HP_ and H5N8_LP_. The dotted line represents the limit of detection for each experiment. Results are representative of an experiment performed three times.

Viral growth kinetics was analyzed in Madin-Darby canine kidney (MDCK) cells, in the DF-1 chicken fibroblast cell line and in the CCL-141 duck fibroblast cell line infected at a multiplicity of infection (MOI) of 10^−5^ tissue culture infectious dose 50 (TCID_50_). H5N8_HP_ replicated with similar kinetics in the absence or presence of exogenous trypsin, demonstrating that proteolytic processing of the HA MBCS was independent of trypsin-like proteases, as expected for a HPAIV (Fig. 1B). By contrast, H5N8_LP_ replication was severely impaired in the absence of trypsin, while its replication kinetics was indistinguishable from that of the H5N8_HP_ in the presence of trypsin (Fig. 1B). These results indicate that mutations introduced in the H5N8_LP_ HA have no negative impact on viral replication kinetics in cell culture beyond the expected trypsin-like proteases requirement for H5N8_LP_ HA proteolytic processing, as expected for a LPAIV.

### H5N8_HP_ and H5N8_LP_ viruses co-infections *in ovo*

Next, we evaluated H5N8_HP_ and H5N8_LP_ replication in embryonated chicken (*Gallus gallus*) and Pekin duck (*Anas platyrhynchos domesticus*) eggs, as they have a complex cellular architecture and provide an intermediate model between monolayer cell culture and *in vivo* studies. To investigate how the host species modulated the interaction between a HPAIV and a LPAIV, we inoculated embryonated chicken and duck eggs via the allantoic cavity either with 10^2^ egg infectious dose 50 (EID_50_) H5N8_HP_ alone, with increasing doses of H5N8_LP_ alone, or with 10^2^ EID_50_ H5N8_HP_ in combination with increasing doses of H5N8_LP_ (Fig. 2A). We inoculated chicken eggs at day 10 and duck eggs at day 11 in order to work at similar development stages (Li et al., 2019). After 24 hours of incubation at 37°C, we harvested the allantoic fluid and quantified the level of virus replication by RT-qPCR using primers specific for H5N8_HP_ or H5N8_LP_ HA. In chicken eggs, we detected equivalent levels of viral RNA following inoculation with 10^2^EID_50_ H5N8_HP_ alone or with H5N8_LP_ alone regardless of the inoculum dose (Fig. 2B). Upon co-infection with H5N8_HP_ and H5N8_LP_ in chicken eggs, the level of H5N8_HP_ and H5N8_LP_-specific viral RNA did not differ from mono-infections regardless of the quantity of H5N8_LP_ (Fig. 2B). Thus, the replication of H5N8_HP_ is not affected by H5N8_LP_, regardless of the amount of H5N8_LP_ co-inoculated in chicken eggs. In duck eggs, we detected similar levels of viral RNA following inoculation with 10^2^ EID_50_ H5N8_HP_ alone or with 10^2^ EID_50_ H5N8_LP_ alone (Fig. 2C). When duck eggs were infected simultaneously with H5N8_HP_ and H5N8_LP_, we observed a decrease of H5N8_HP_ viral RNA levels that correlated with the amount of H5N8_LP_ co-inoculated, indicating that the H5N8_LP_ antagonized H5N8_HP_ replication in duck eggs (Fig. 2C). The *in ovo* experiments provide evidence that a ≥100-fold excess of H5N8_LP_ significantly antagonized H5N8_HP_ replication in duck eggs, while H5N8_LP_ did not affect H5N8_HP_ replication in chicken eggs regardless of the amount of H5N8_LP_. This observation thus indicates that the H5N8_HP_ had a stronger selective advantage over the H5N8_LP_ in chicken embryonated eggs than in duck embryonated eggs.

**Fig. 2.**
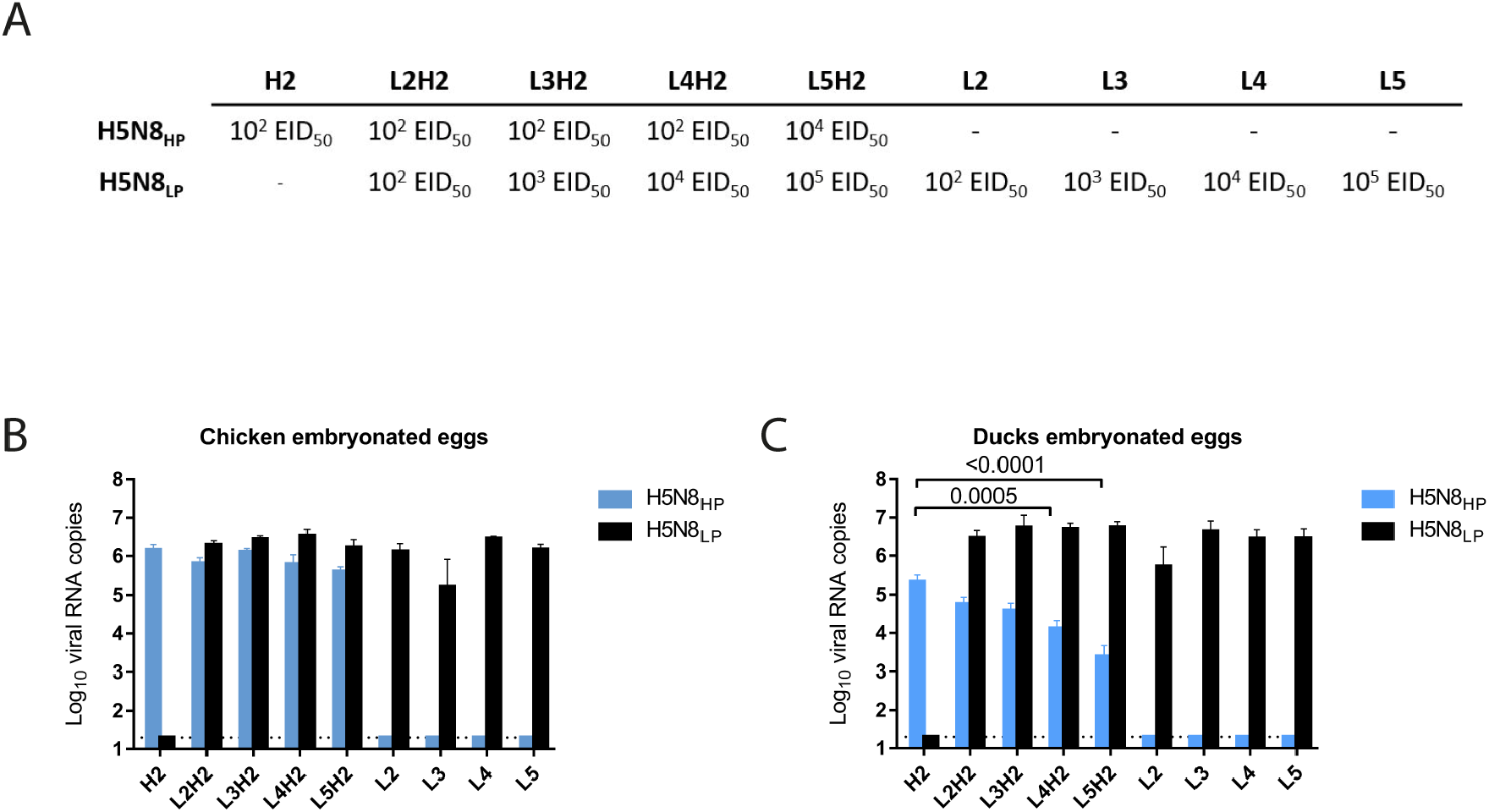
H5N8_HP_ has a stronger advantage over H5N8_LP_ in embryonated chicken eggs than in embryonated duck eggs. **(A)** EID_50_ doses used in the *in ovo* experiments. **(B-C)** Chicken **(B)** and duck **(C)** embryonated eggs were infected in the allantoic cavity either with H5N8_HP_ or H5N8_LP_ alone, or with a combination of both. Following a 24 hours incubation at 37°C, viral RNA was extracted from the allantoic fluid and HA levels were determined by RT-qPCR using H5N8_HP_ or H5N8_LP_ specific primers. Statistical analysis: chicken embryonated eggs: n=9 (experiment performed twice); duck embryonated eggs: n=26 (experiment performed three times); one-way ANOVA with Tukey’s multiple comparisons test. Results are expressed as means ± SEM. The dotted line represents the limit of detection for each experiment.

### Potentiation of HPAIV replication and pathogenesis by LPAIV in chickens

In a preliminary experiment (data not shown), we observed transient low-level oropharyngeal viral RNA shedding in 6/8 of the 10^6^ EID_50_ H5N8_LP_ infected chickens (*Gallus gallus*), whereas all chickens infected with 10^7^ EID^50^ H5N8^LP^ shed high levels of viral RNA for a prolonged time. No clinical signs were observed in chickens inoculated with H5N8^LP^. Furthermore, by using increasing doses of H5N8^HP^, we evaluated the H5N8^HP^ chicken infectious dose 50 at 6×10^3^ EID_50_ and noticed that mortality reached 100% within four days with doses higher than 10^5^ EID_50_. H5N8_HP_-infected sick birds presented dyspnea that quickly progressed to severe dyspnea, associated with anorexia and lethargy. These results provided evidence that the H5N8_HP_ and H5N8_LP_ caused, respectively, commonly observed HPAIV and LPAIV-associated infection patterns in chickens (EFSA Panel on Animal Health and Welfare (AHAW) et al., 2017).

Based on these results, we investigated the interaction between a HPAIV and a LPAIV in chickens. Four-week old chickens were assigned one of six groups: L7, H3, H4, L7H3, L7H4 or NI (Fig. 3A). L7 animals were infected with 10^7^ EID_50_ H5N8_LP_. H3 animals were infected with 10^3^ EID_50_ H5N8_HP_. H4 animals were infected with 10^4^ EID_50_ H5N8_HP_. L7H3 animals were infected with a mixture of 10^7^ EID_50_ H5N8_LP_ and 10^3^ EID_50_ H5N8_HP_. L7H4 animals were infected with a mixture of 10^7^ EID^50^ H5N8^LP^ and 10^4^ EID_50_ H5N8_HP_. Finally, non-infected animals from group NI were administered vehicle only. Neither mortality nor clinical signs were observed in chickens from group L7 and group H3 (Fig. 3B). Chickens in the H4 group reached 45% mortality (Fig. 3B). When chickens were infected with a mixture of H5N8_LP_ and H5N8_HP_, we observed an increase in mortality compared to H5N8HP mono-infected chickens: mortality reached 18% in the L7H3 group, compared to 0% in the H3 group; and 72% in the L7H4 group, compared to 45% in the H4 group (Fig. 3B). Although the increase in mortality did not reach statistical significance, these results indicate that H5N8_LP_ and H5N8_HP_ synergized, resulting in increased pathogenesis in co-infected chickens compared to H5N8_HP_ mono-infected chickens.

**Fig. 3.**
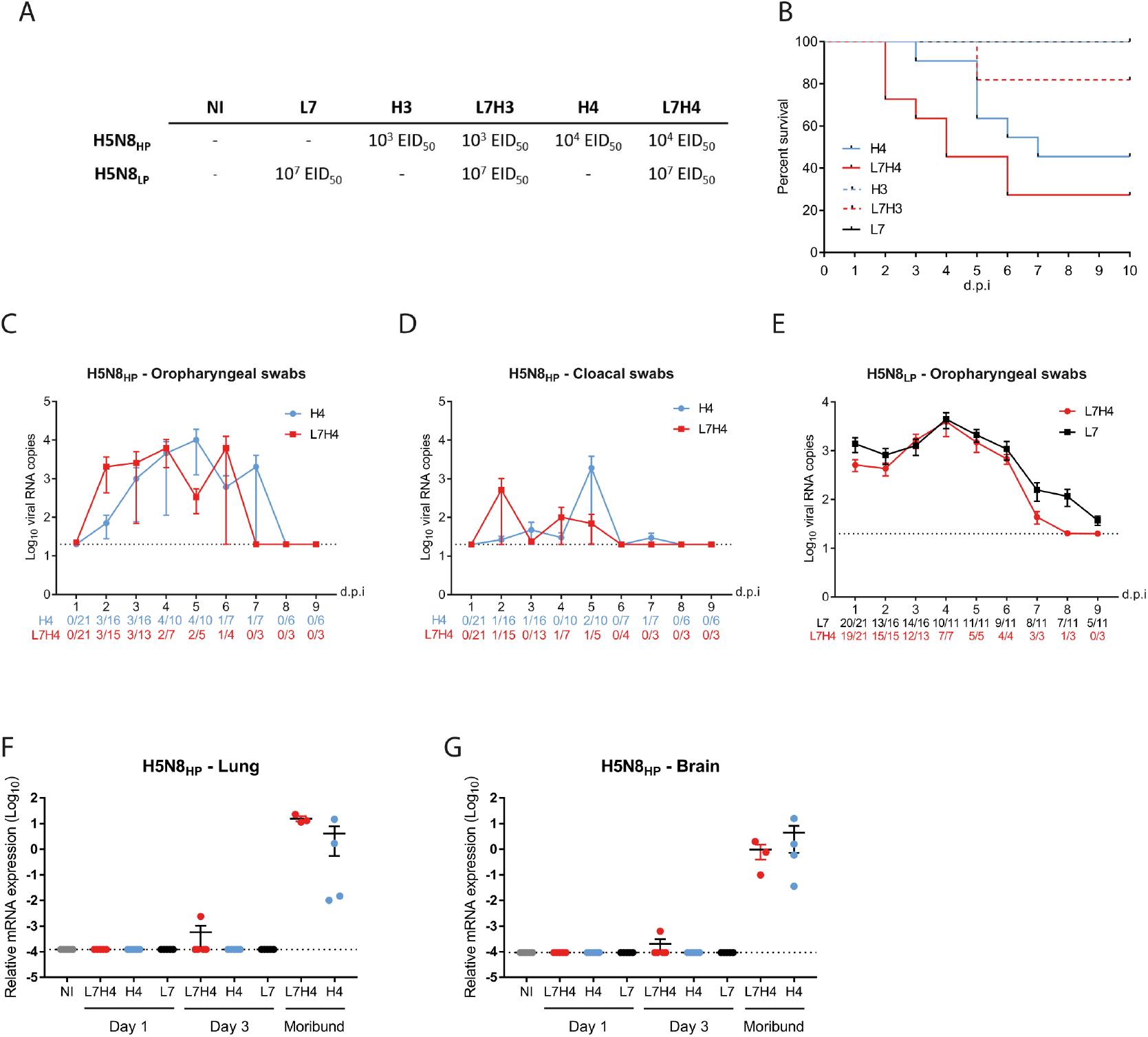
H5N8_HP_ and H5N8_LP_ coinfection in chickens results in increased pathogenesis. Chickens were inoculated via the choanal route with H5N8_LP_ alone (L7), H5N8_HP_ alone (H3 and H4) or with a combination of H5N8_LP_ and H5N8_HP_ (L7H3 and L7H4). **(A)** EID_50_ doses used in the *in vivo* chicken experiments. **(B)** Survival curves. Statistical analysis: Log-rank (Mantel-Cox) test. **(C-E)** Viral shedding was analyzed by quantifying HA RNA levels by RT-qPCR using H5N8_HP_ specific primers from RNA extracted from oropharyngeal swabs **(C)** and cloacal swabs **(D)** or by RT-qPCR using H5N8_LP_ specific primers from RNA extracted from oropharyngeal swabs **(E)**. Numbers of H5N8_HP_ or H5N8_HP_ swab-positive animals are indicated below each time point. Statistical analysis: two-tailed Mann-Whitney test. Results are expressed as means ± SEM. The dotted line represents the limit of detection. **(F-G)** H5N8_HP_ load was analyzed from total RNA extracted from the lungs **(F)** and the brain **(G)** using H5N8_HP_ specific primers. HA RNA levels were normalized using the 2^-ΔCt^ method. Statistical analysis: one-way ANOVA with Tukey’s multiple comparisons test. Results are expressed as means ± SEM. The dotted line represents the limit of detection. dpi, days post-infection.

**Fig. 4.**
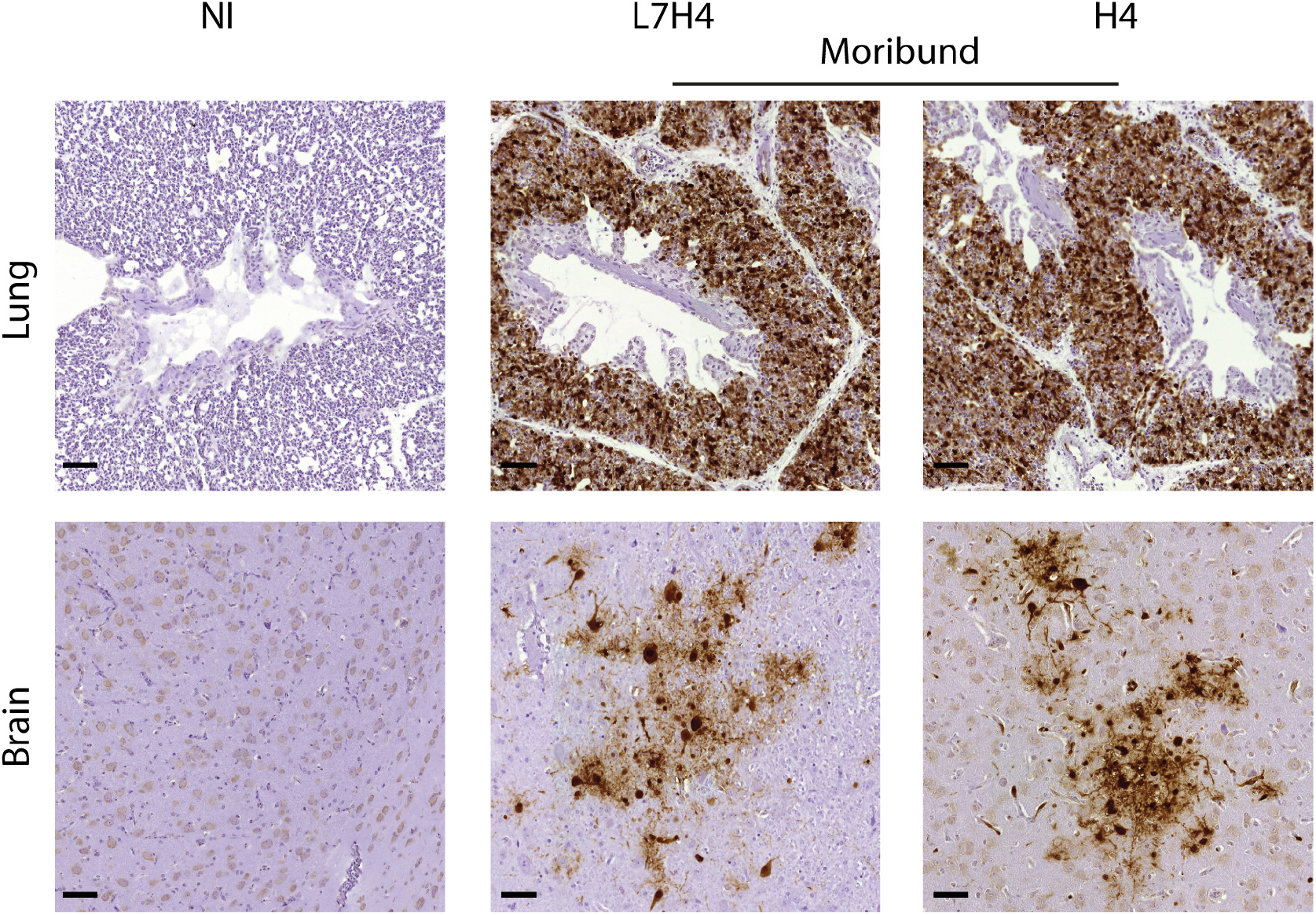
Immunohistochemical anti-NP staining of hematoxylin-counterstained chickens lung or brain sections. Analysis was performed on non-infected (NI) and moribund chickens. Scale bar, 50 µm.

**Fig. 5.**
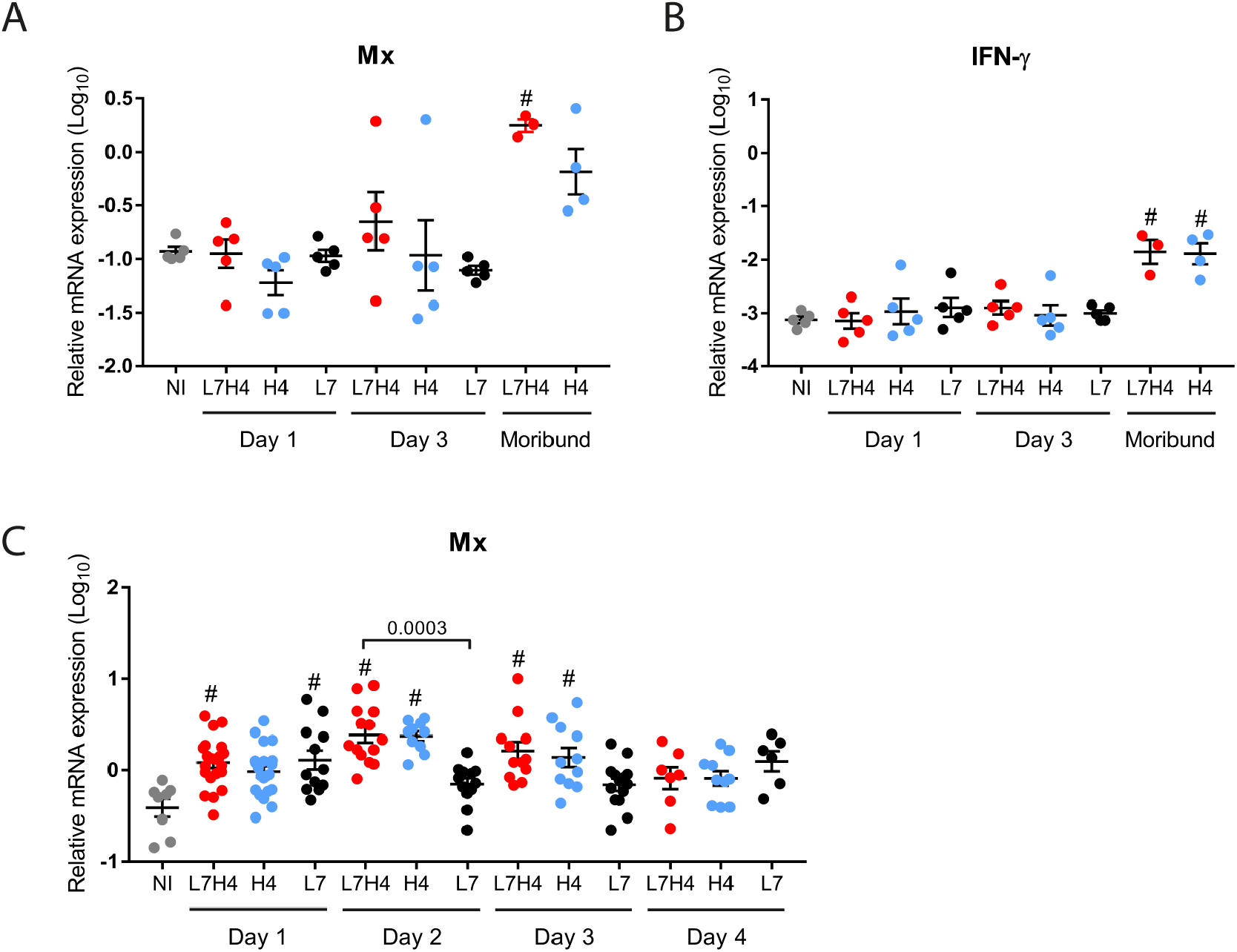
Mx and IFN-γ mRNA expression following chickens infection. **(A-B)** mRNA expression levels of Mx **(A)** and IFN-γ **(B)** determined by RT-qPCR performed on lung total RNA. **(C)** mRNA expression levels of Mx determined by RT-qPCR performed on RNA extracted from oropharyngeal swabs. mRNA levels were normalized using the 2^-ΔCt^ method. Statistical analysis: one-way ANOVA with Tukey’s multiple comparisons test. Results are expressed as means ± SEM. #, p < 0.05 compared to non-infected (NI) animals.

**Fig. 6.**
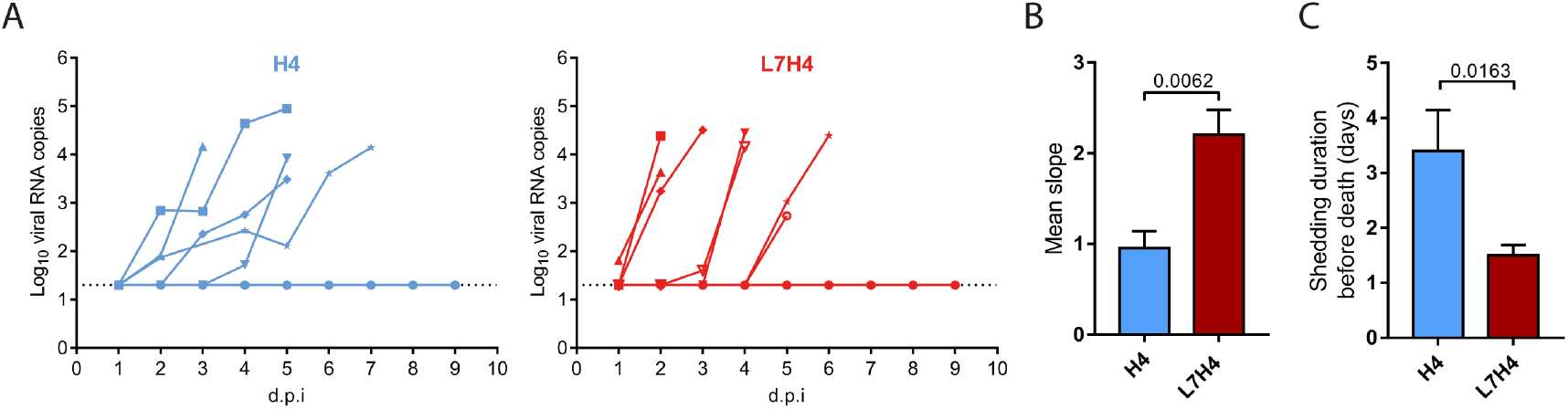
H5N8_HP_ and H5N8_LP_ coinfection in chickens results in higher H5N8_HP_ growth. **(A)** Individual H5N8_HP_ oropharyngeal shedding curves in H4 chickens (blue lines) and L7H4 chickens (red lines). Each curve corresponds to a single animal from the onset of H5N8_HP_ excretion to death. **(B)** The slopes of individual H5N8_HP_ excretion curves were calculated. **(C)** H5N8_HP_ shedding duration: number of days where chickens were H5N8_HP_ oropharyngeal swab-positive. Statistical analysis: two-tailed Mann-Whitney test. Results are expressed as means ± SEM. The dotted line corresponds to the limit of detection. dpi, days post-infection.

We measured viral shedding from oropharyngeal and cloacal swabs by quantifying viral RNA using RT-qPCR with primers detecting specifically H5N8_LP_ or H5N8_HP_ HA sequences. Regardless of the experimental group, all H5N8_HP_ swab-positive chickens eventually developed severe dyspnea and either died or were euthanized when they reached humane endpoints, as previously observed (Leyson et al., 2019). In line with the mortality rate (Fig 3B), no H5N8_HP_ swab-positive chicken was found in the H3 group, while only two chickens tested positive for H5N8_HP_ in the L7H3 group. Because of the low rate of infection in the H3 and L7H3 groups, these animals were excluded from further analyses. H5N8_HP_ oropharyngeal and cloacal shedding was higher in group L7H4 compared to group H4 at 2 days post-infection (dpi), but the difference was not statistically significant (Fig. 3C&D). Average H5N8_HP_ oropharyngeal and cloacal shedding remained otherwise similar between the L7H4 and the H4 groups and was not associated with different H5N8_HP_ transmission rates to contact chickens introduced in the poultry isolators at 1 dpi (data not shown). H5N8_LP_ oropharyngeal shedding did not differ between the L7 and L7H4 group (Fig. 3E).

### Antagonism of HPAIV replication and pathogenesis by LPAIV in ducks

We inoculated groups of 4-week old Pekin ducks (*Anas platyrhynchos domesticus*) with H5N8_HP_ alone, H5N8_LP_ alone, or a mixture containing both H5N8_LP_ and H5N8_HP_. Ducks were assigned one of four groups: L7, H4, L7H4 or NI (Fig. 7A). L7 animals were infected with 10^7^ EID_50_ H5N8_LP_. H4 animals were infected with 10^4^ EID_50_ H5N8_HP_. L7H4 animals were infected with a mixture of 10^7^ EID_50_ H5N8_LP_ and 10^4^ EID_50_ H5N8_HP_. Non-infected animals from group NI were administered vehicle only. Neither mortality nor clinical signs were observed in any duck of group L7 (Fig. 7B). Mortality reached 87% in the H4 group and was preceded by predominantly neurological clinical signs, in accordance with the pronounced neurotropism of clade 2.3.4.4 group B H5N8 viruses in ducks (Grund et al., 2018; Kleyheeg et al., 2017; More et al., 2017). Mortality was significantly reduced to 37% in the L7H4 group. Thus, H5N8_LP_ antagonized H5N8_HP_ pathogenesis in co-infected ducks, in contrast to what we observed in chickens.

**Fig. 7.**
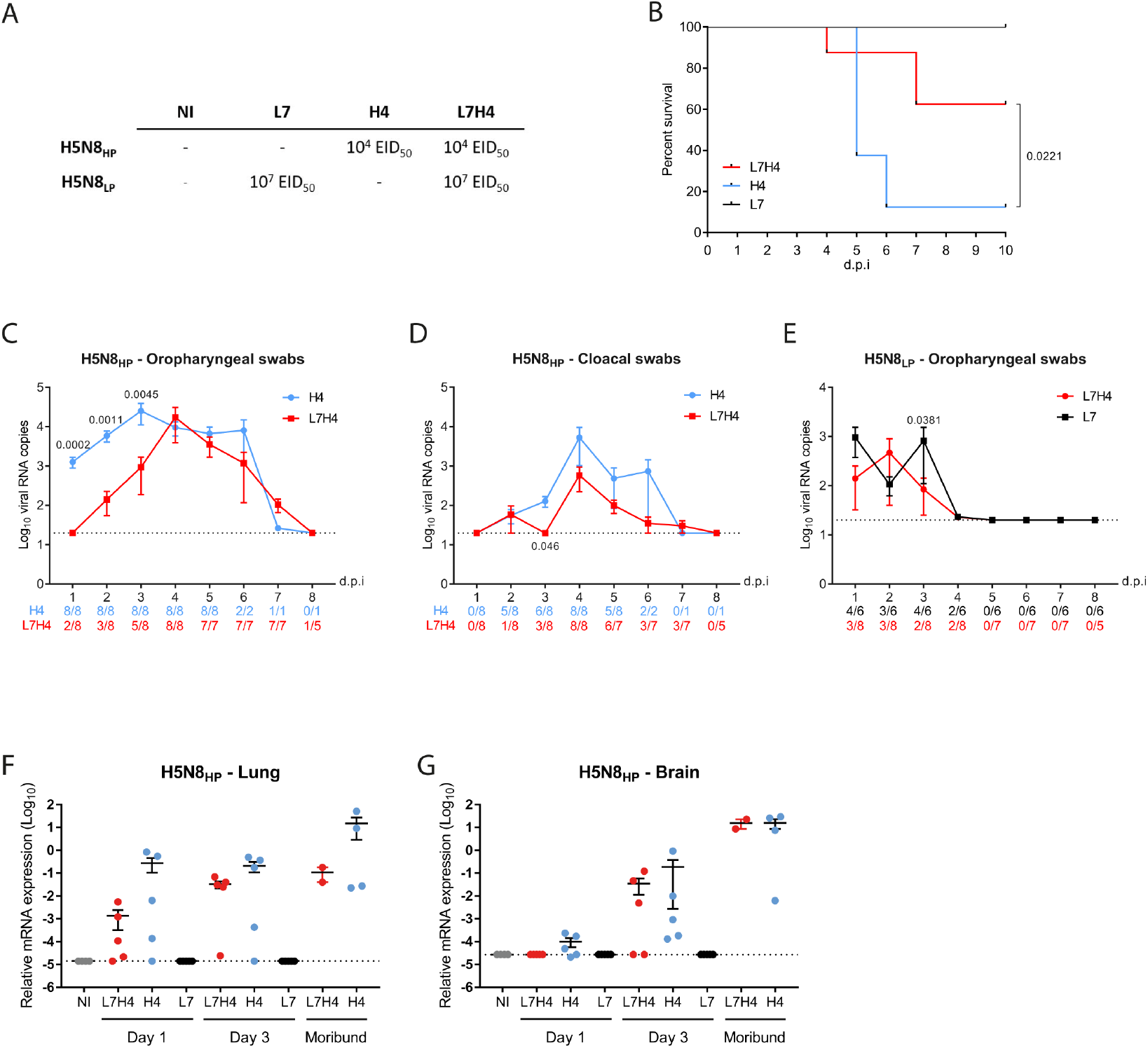
H5N8_HP_ and H5N8_LP_ coinfection in ducks results in decreased pathogenesis. Ducks were inoculated via the choanal route with H5N8_LP_ alone (L7), H5N8_HP_ alone (H4) or with a combination of H5N8_LP_ and H5N8_HP_ (L7H4). **(A)** EID_50_ doses used in the *in vivo* duck experiments. **(B)** Survival curves. Statistical analysis: Log-rank (Mantel-Cox) test. **(C-E)** Viral shedding was analyzed by quantifying HA RNA levels by RT-qPCR using H5N8_HP_ specific primers from RNA extracted from oropharyngeal swabs **(B)** and cloacal swabs **(D)** or by RT-qPCR using H5N8_LP_ specific primers from RNA extracted from oropharyngeal swabs **(E)**. Numbers of H5N8_HP_ or H5N8_HP_ swab-positive animals are indicated below each time point. Statistical analysis: two-tailed Mann-Whitney test. Results are expressed as means ± SEM. The dotted line represents the limit of detection. **(F-G)** H5N8_HP_ load was analyzed from total RNA extracted from the lungs **(F)** and the brain **(G)** using H5N8_HP_ specific primers. HA RNA levels were normalized using the 2^-ΔCt^ method. Statistical analysis: one-way ANOVA with Tukey’s multiple comparisons test. Results are expressed as means ± SEM. The dotted line represents the limit of detection. dpi, days post-infection.

To determine how this related to the levels of virus replication, we analyzed H5N8_LP_ and H5N8_HP_ viral RNA shedding from oropharyngeal and cloacal swabs using RT-qPCR. Oropharyngeal H5N8_HP_ shedding was significantly reduced in group L7H4 compared to group H4 in the first days post-infection (dpi), with a hundred-fold difference at 1 dpi (p<0.001) and a ten-fold difference at 2 and 3 dpi (p<0.01) (Fig. 7C). Cloacal H5N8_HP_ shedding was also decreased in group L7H4, but the difference only reached statistical difference at day 3, with a ten-fold reduction (p<0.05) (Fig. 7D). There was no difference in H5N8_LP_ shedding between groups L7H4 and L7 (Fig. 7E).

Next, we analyzed viral RNA levels in the lungs and brain. H5N8_LP_ was not detected in the lungs or brain, indicating that H5N8_LP_ replicated mostly in the upper respiratory tract of ducks. In contrast to chickens, we detected H5N8_HP_ nucleic viral RNA from the lungs and brain of most animals at 1 and 3 dpi (Fig. 7F&G). H5N8_HP_ viral RNA load was decreased in the lungs and brain of H5N8_LP_ and H5N8_HP_ co-infected birds (group L7H4), compared to H5N8_HP_ mono-infected birds (group H4). High levels of H5N8_HP_ viral RNA were also detected in the lungs and brain of moribund ducks, which developed predominantly neurological signs.

Extensive viral antigen staining was detected by immunohistochemistry in the brain of moribund animals, while more modest viral antigen staining was observed in the lungs, with no difference between groups L7H4 and H4 (Fig. 8A&B). We then evaluated the expression of host immune response markers in the lungs and brain using RT-qPCR. Mx mRNA expression was significantly increased in the lungs of infected ducks belonging to the L7H4 and H4 groups at 1 and 3 dpi, as well as in moribund animals (Fig. 9A). Compared to NI duck, we also observed an upregulation of Mx mRNA expression in the lungs of H5N8_LP_ mono-infected ducks (L7 group) at 1 and 3 dpi, although the difference did not reach statistical significance. IFN-γ mRNAs levels were significantly higher in the lungs of H5N8_HP_ mono-infected ducks (group H4) at 1 dpi compared to NI, L7H4 and L7 ducks (Fig. 9B). Similar results were observed in the brain (data not shown). To evaluate the antiviral innate immune response in the upper respiratory tract, we measured Mx mRNA expression by RT-qPCR from RNA extracted from oropharyngeal swabs. Compared to NI ducks, Mx mRNA was increased in all infected groups (Fig. 9C). In contrast to chickens, Mx mRNA expression level remained high in H5N8_LP_ mono-infected ducks from the L7 group (Fig. 9C).

**Fig. 8.**
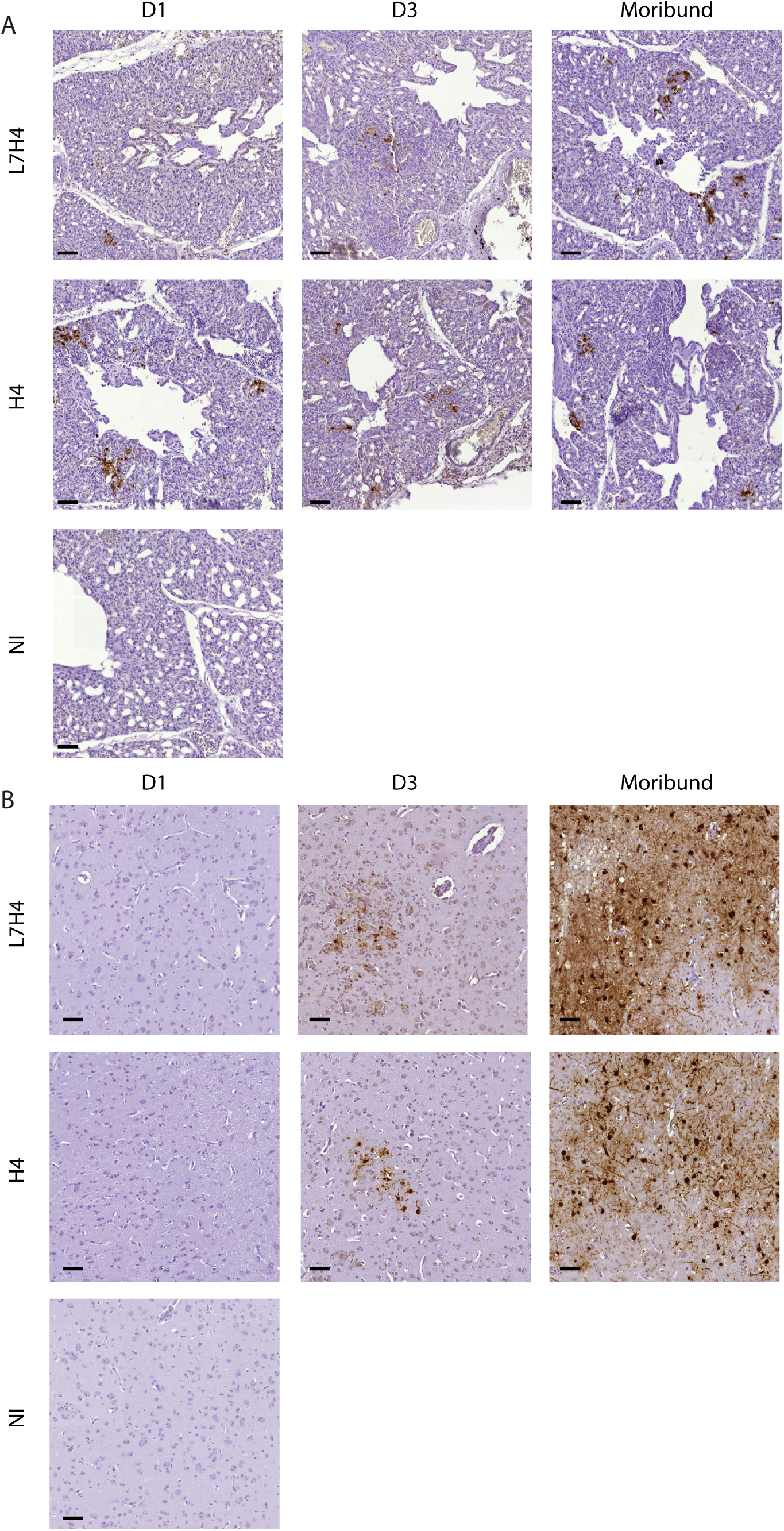
Immunohistochemical anti-NP staining of hematoxylin-counterstained ducks lung or brain sections. Analysis was performed on ducks necropsied at 1 (D1) and 3 days (D3) post infection, moribund ducks and non-infected (NI) ducks. **(A)** Lung. **(B)** Brain. Scale bar, 50 µm.

**Fig. 9.**
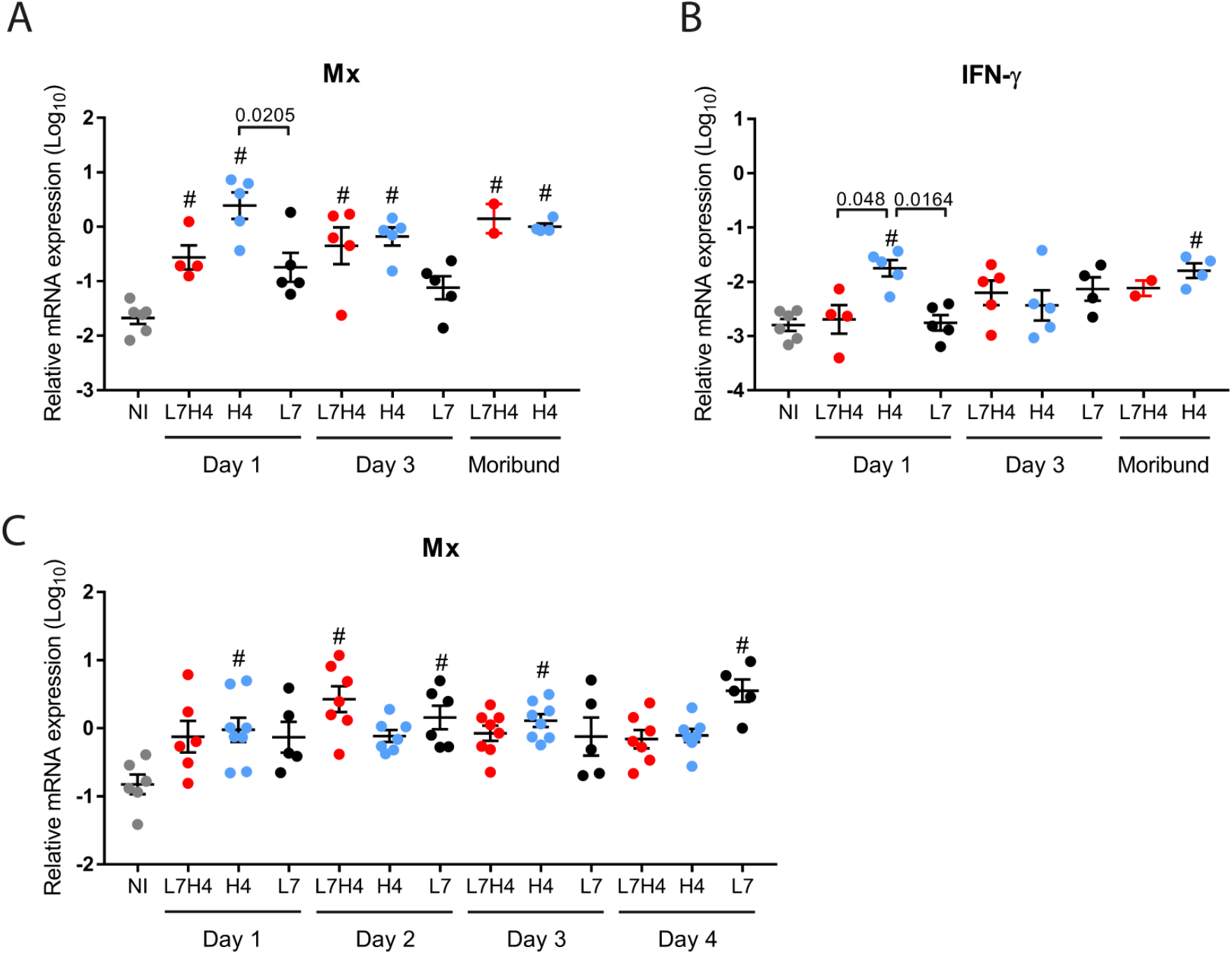
Mx and IFN-γ mRNA expression following infection in ducks. **(A-B)** mRNA expression levels of Mx **(A)** and IFN-γ **(B)** determined by RT-qPCR performed on lung total RNA. **(C)** mRNA expression levels of Mx determined by RT-qPCR performed on RNA extracted from oropharyngeal swabs. mRNA levels were normalized using the 2^-ΔCt^ method. Statistical analysis: one-way ANOVA with Tukey’s multiple comparisons test. Results are expressed as means ± SEM. #, p < 0.05 compared to non-infected (NI) animals.

To further compare the antiviral innate immune response in ducks and chickens, we plotted the levels of Mx mRNA as a function of the level of viral RNA in the lungs (Fig. 10A). We observed a correlation between the level of Mx mRNA and the level of viral RNA in ducks (Pearson r=0.64; p<0.001), confirming that the intensity of the antiviral innate immune response is proportional to the level of viral RNA, as previously reported (Baccam et al., 2006; Hagenaars et al., 2016; Soubies et al., 2013; Volmer et al., 2011). This correlation was less visible in chickens, possibly because viral RNA was only found in the lungs of a limited number of chickens (Pearson r=0.50; p=0.21). The Mx/HA ratios were distributed to higher values in ducks compared to chickens (Fig. 10A), resulting in mean Mx/HA ratios that were significantly higher in ducks than in chickens (Fig. 10B). This result demonstrates that ducks mounted a more potent antiviral innate immune response against influenza virus infection than chickens.

**Fig. 10.**
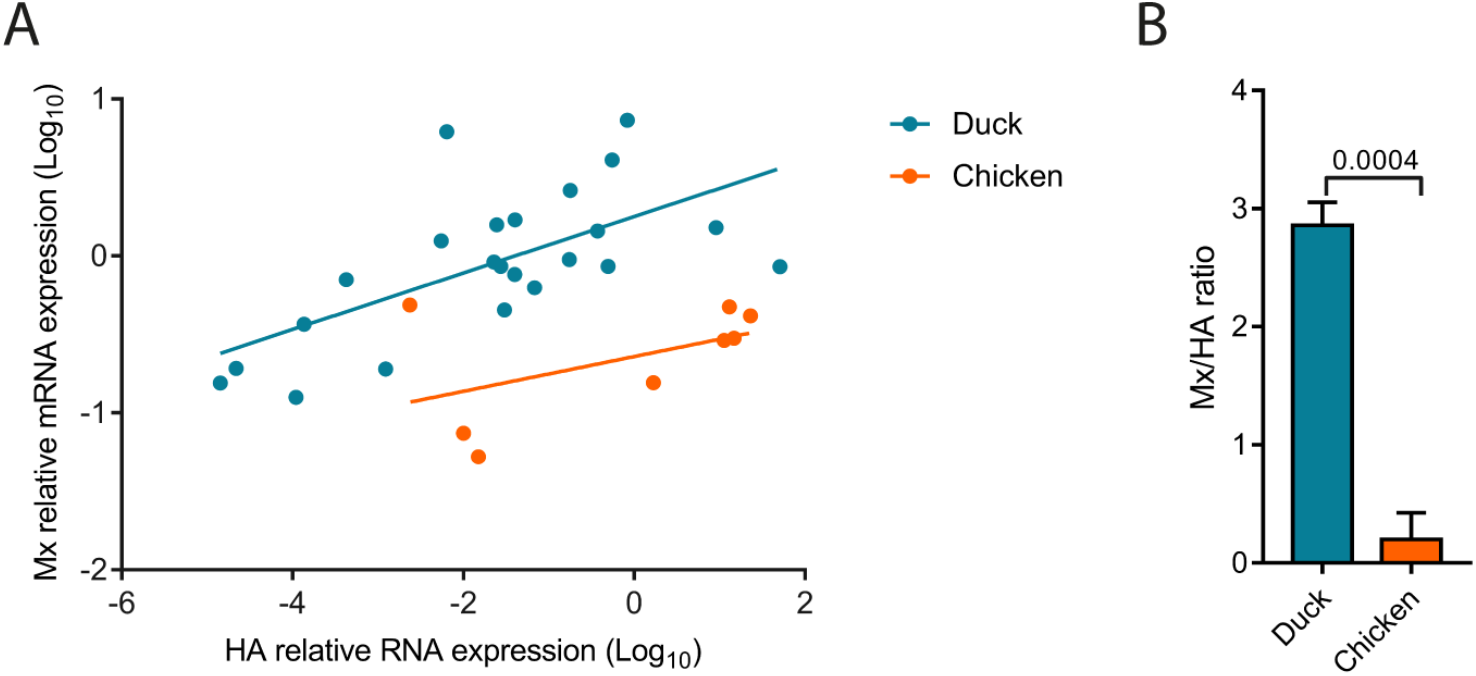
Ducks mount a more potent antiviral innate immune response compared to chickens. **(A)** Correlation between Mx mRNA levels and viral RNA levels in the lungs. **(B)** Mean Mx/HA ratios. Mx mRNA and HA RNA levels were normalized using the 2^-ΔCt^ method. Statistical analysis: two-tailed Mann-Whitney test. Results are expressed as means ± SEM.

## DISCUSSION

To model the intra-host competition between a newly formed HPAIV and its parental LPAIV, we inoculated chickens and ducks with an H5N8_HP_ virus and an H5N8_LP_ virus differing solely at the level of the HA cleavage site. When inoculated alone, H5N8_HP_ replication and pathogenesis were equivalent in chickens and ducks. However, when chickens and ducks were co-inoculated with this pair of viruses, we found that the H5N8_HP_ had a stronger selective advantage over the H5N8_LP_ in chickens than in ducks. Surprisingly, we observed that the H5N8_LP_ increased H5N8_HP_ replication and pathogenesis in chickens. By contrast, we observed that the H5N8_LP_ antagonized H5N8_HP_ replication and pathogenesis in ducks. Thus, the nature of the interaction between the HPAIV and the parental LPAIV may be a critical determinant of HPAIV emergence, which we show here to depend on the host species.

The interaction between co-infecting viruses ranges from synergy to antagonism depending on the virus pairs and the host (Andreu-Moreno et al., 2020; Bou et al., 2019; DaPalma et al., 2010; Domingo-Calap et al., 2019; Kumar et al., 2018; Opatowski et al., 2018; Phipps et al., 2020). Functional complementation occurs when one virus provides a function that is absent or less efficient in the other virus. As the genomes of the H5N8_HP_ and H5N8_LP_ differ solely at the level of the HA cleavage site, it is unlikely that the H5N8_LP_ is able to provide a function missing in the H5N8_HP_. We therefore speculate that the H5N8_LP_-mediated increase in H5N8_HP_ replication and pathogenesis observed in chickens could be due to complementation of semi-infectious H5N8_HP_ particles by the H5N8_LP_ inoculated at a higher dosage (Brooke, 2017). Such complementation is likely to occur both in chickens and in ducks. However, this complementation may be undetectable in ducks due to the dominant effect of host-mediated antagonism between the H5N8_LP_ and the H5N8_HP_. The antagonism between the H5N8_LP_ and the H5N8_HP_ is unlikely to be due to viral interference due to receptor or cell machinery usage, as no competition between the H5N8_LP_ and the H5N8_HP_ was observed in chickens, although the H5N8_LP_ replicated to higher levels and persisted longer in chickens than in ducks. In contrast to chickens, ducks have a functional RIG-I receptor, which has been proposed to contribute to the more efficient type I interferon mediated-antiviral innate immune response against influenza viruses observed in ducks compared to chickens (Barber et al., 2010; Burggraaf et al., 2014; Cornelissen et al., 2013, 2012; Kuchipudi et al., 2014; Magor et al., 2013; Soubies et al., 2013). The H5N8_LP_ and the H5N8_HP_ are likely to be intrinsically equally sensitive to the antiviral effects of type I interferon. However, a stronger antiviral innate immune response could impose a stronger selective pressure on the minority variant (Zwart and Elena, 2015). In line with this hypothesis, type I interferon antiviral innate immune response was shown to constitute a strong bottleneck shaping poliovirus population in mice (Kuss et al., 2008). Similarly, treatment of hepatitis C virus-infected patients with type I IFN resulted in a dramatic reduction of viral diversity (Farci et al., 2002). Furthermore, a delayed type I IFN response in influenza A virus-infected obese mice promoted minority variant emergence, while type I interferon treatment reduced viral diversity (Honce et al., 2020). Altogether, these results indicate that the innate immunity can shape viral populations. By decreasing viral replication and increasing the within-host competition between variants, a strong innate immune response may inhibit the selection of variants that arise through *de novo* mutations during infection of an individual, unless these variants have a strong selective advantage within this individual. We suggest that the more pronounced antiviral innate immune response observed in ducks compared to chickens may impact the H5N8_HP_ minority variant more severely in ducks than in chickens. Admittedly, the differential outcome of H5N8_HP_ and H5N8_LP_ co-infections in chickens and ducks may also depend on other host factors, including microbiota composition or receptor distribution (Figueroa et al., 2020; Hiono et al., 2016).

Recent work investigating the interaction between an H7N7 HPAIV and a closely related H7N7 LPAIV field isolate demonstrated that the LPAIV inhibited HPAIV replication in chickens when the LPAIV was inoculated at a ≥ 100-fold higher dose (Graaf et al., 2018). These results indicate that the nature of the interaction between the HPAIV and the LPAIV may depend on the viral strain. If competition between a LPAIV and a HPAIV occurred in chickens, we speculate that the competition could even be stronger in ducks, because of the stronger antiviral innate immune-mediated selective pressure imposed on the minority HPAIV. However, further work is needed to properly address this question. These studies should ideally be performed using different pairs of reverse-genetics engineered HPAIV and LPAIV differing solely at the level of the HA cleavage site to specifically investigate how the selective advantage conferred by the MBCS depends on the host species.

Epidemiological investigations of HPAIV outbreaks indicate that the vast majority of HPAIV emerge upon replication and inter-individual transmission of a LPAIV in *Galliformes*, such as chickens and turkeys, although *Anseriformes*, such as ducks and geese, are considered as the main reservoirs for LPAIV (Abdelwhab et al., 2013; EFSA Panel on Animal Health and Welfare (AHAW) et al., 2017; Lee et al., 2021; Richard et al., 2017). In this study, we focused on the two main representatives of the *Galliformes* and *Anseriformes* orders: chickens, which are with current global production of 22 billion chickens the main source of animal protein for human consumption worldwide and Pekin ducks, which are also intensively raised in farms and represent the best characterized *Anseriformes* in respect to the interaction with influenza viruses (Lee et al., 2021; “Poultry species | Gateway to poultry production and products | Food and Agriculture Organization of the United Nations,” n.d.). How the LPAIV and the newly formed HPAIV would interact in other bird species, such as turkeys, or in wild dabbling or diving ducks, which have a prominent role in the ecology of avian influenza viruses will be the subject of future studies (Bodewes and Kuiken, 2018; EFSA Panel on Animal Health and Welfare (AHAW) et al., 2017; Lee et al., 2021). In addition to host factors, anthropic factors are also likely to play a major role in the emergence of HPAIV and may contribute to the higher frequency of HPAIV emergence observed in chickens that are usually raised at higher densities than ducks (Capua and Marangon, 2000; Dhingra et al., 2018; M.c et al., 2009; Richard et al., 2017). Estimating to which extent farming processes contribute to the emergence of HPAIV is however a difficult task and therefore their contribution to HPAIV emergence remains currently unclear.

To minimize the risk of HPAIV evolution, present European regulation and national regulations in many countries require the culling of all birds in flocks infected with H5 and H7 LPAIV (EFSA Panel on Animal Health and Welfare (AHAW) et al., 2017). Culling of H5 or H7 LPAIV infected flocks aims at eliminating LPAIV from the susceptible poultry population before they have a chance to evolve into HPAIV, and is therefore considered a preventive biosecurity measure. However, the broader public, breeders and professional breeder societies question the need for preventive culling for economical, animal welfare and breeder welfare issues (Organization, 2008). Indeed, preventive culling of animals, especially healthy animals, which is commonly the case in LPAIV infected *Anseriformes*, is poorly accepted. Identifying how host factors modulate HPAIV emergence therefore provides important scientific data that may be taken into account by animal health policy makers in the future.

## MATERIALS AND METHODS

### Viruses

The eight segments from the HPAIV field isolate A/mulard duck/France/171201g/2017 (H5N8) were cloned in a pHW2000 plasmid vector using a set of universal primers in order to generate a reverse-genetics engineered wild-type HPAIV H5N8 (H5N8_HP_) (Hoffmann et al., 2001, 2000). Using site-directed mutagenesis, a 9 nucleotides deletion was performed on the HA cleavage site sequence, along with two single nucleotide polymorphism on both sides of the deletion to obtain a reverse-genetics engineered LPAIV H5N8 (H5N8_LP_) according to the manufacturer’s instructions (In-Fusion, Takara Bio, France). 2.5×10^5^ HEK 293T cells cultured in 6-well plates were transfected in Opti-MEM medium, using the lipophilic transfection reagent LTX with Plus reagent (Invitrogen, Thermo Fisher Scientific Inc., Canada), either with 0.5µg of each seven common H5N8_HP/LP_ pHW2000 plasmids (PB2, PB1, PA, NA, NP, M and NS) and with 0.5µg of HA_HP_ or HA_LP_. L-(tosylamido-2-phenyl)ethyl chloromethyl ketone (TPCK) treated trypsin (TPCK-treated) was added 24 hours post-transfection at a 0.5µg/ml final concentration. 48 hours post-transfection, scraped cells and culture medium were transferred on MDCK cells grown with Opti-MEM supplemented with 0.5µg/ml TPCK-treated trypsin. 72 hours later, cell supernatant was collected. To produce viral stock, both viruses were then propagated in 10-day-old chicken specific-pathogen-free (SPF) embryonated chicken eggs (INRAE, PFIE, Nouzilly, France), by inoculation in the allantoic cavity of 100µL of 1:100 dilutions of infected MDCK cells supernatant. Infectious allantoic fluid was harvested at 72 hours postinoculation and titrated in 10-day-old SPF embryonated chicken eggs to determine the 50% egg infective dose (EID_50_)/mL using the Reed-Muench method. The identity of amplified viruses was verified by Sanger sequencing of each viral gene segment (H5N8_HP_: accession numbers MK859904 to MK859911; H5N8_LP_: accession numbers MK859926 to MK859933). These viruses were referenced by the French biotechnology ethics committee (Haut Conseil des Biotechnologies) and were manipulated exclusively in biosafety level 3 laboratories.

### *In vitro* infections

MDCK cells, DF-1 cells and CCL-141 cells were cultured in DMEM containing 1% antibiotics (penicillin/streptomycin) and 10% fetal bovine serum. When cells reach 90% confluency, they were washed with PBS and infected either with H5N8_HP_ or with H5N8_LP_ at a multiplicity of infection (MOI) of 10^−5^ TCID_50_. After one hour, the inoculum was removed, cells were washed twice with PBS and Opti-MEM supplemented with 0.5µg/mL TPCK-treated trypsin (Pierce, Thermo Fisher Scientific Inc., Canada) was added. Viral RNA extraction was performed on 140µL supernatant collected at 1, 24, 48 and 72 hours post-infection according to the manufacturer’s instructions (QIAamp viral RNA; Qiagen, Toronto, Canada). Influenza nucleic acid load was determined by RT-qPCR using primers targeting both H5N8_LP_ and H5N8_HP_ HA gene in a final volume of 10µL. Mixes were prepared according to the manufacturer’s instructions (iTaq Universal SYBR green One-Step kit, BioRad) with 1µL of RNA and a final concentration of 0.3µM of each primer.

### *In ovo* co-infection experiments

Specific pathogen-free (SPF) White Leghorn (PA12) embryonated chicken eggs (PFIE, INRAE, Nouzilly, France) and Pekin Duck (ST5 Heavy) embryonated eggs (ORVIA-Couvoir de la Seigneurtière, Vieillevigne, France) were respectively incubated for 10 and 11 days at 37°C to work at similar stages of development (Li et al., 2019). They were then inoculated in the allantoic cavity with 200µL virus diluted in PBS containing 1% antibiotics (penicillin/streptomycin). Eggs from group H2 were infected with 10^2^ EID_50_ H5N8_HP_. Eggs from groups L2, L3, L4 and L5 were infected with 10^2^ EID_50,_ 10^3^ EID_50,_ 10^4^ EID_50_, and 10^5^ EID_50_ H5N8_LP_ respectively. Eggs from groups L2H2, L3H2, L4H2 and L5H2 were infected with a mixture of 10^2^ EID_50_ H5N8_HP_ and increasing doses of H5N8_LP_: 10^2^, 10^3^, 10^4^ and 10^5^ EID_50_ H5N8_LP_ respectively. Each group contained 9 to 29 eggs from two to three independent experiments. 24 hours post-infection eggs were incubated at 4°C overnight and allantoic fluid was harvested for RNA extraction.

### Animals and groups

Chickens and ducks experiment were conducted separately. One-day-old Pekin ducklings (*Anas platyrhyncos domesticus*, ST5 heavy) were obtained from a commercial hatchery (ORVIA-Couvoir de la Seigneurtière, Vieillevigne, France) and one-day-old White Leghorn chicks (*Gallus gallus domesticus*, PA12) from a research hatchery (PFIE, INRAE, Nouzilly, France). Animals were fed *ad libitum* with a starter diet and housed in biosafety level II facilities for 2 weeks in a litter-covered floor pen at the National Veterinary School of Toulouse, France. They were then transferred into a biosafety level III facility, equipped with bird isolators (I-Box; Noroit, Nantes, France) ventilated under negative pressure with HEPA-filtered air. Chicken preliminary experiment: 39 chickens were randomly assigned to four groups: 10 animals were assigned to group H3 and were infected with 10^3^ EID_50_ H5N8_HP_. 10 animals were assigned to group H4 and were infected with 10^4^ EID_50_ H5N8_HP_. 8 animals were assigned to group L6 and were infected with 10^6^ EID_50_ H5N8_LP_. 11 animals were assigned to group L7 and were infected with 10^7^ EID_50_ H5N8_LP_.

Chickens co-infection experiment: A total of 130 chickens were randomly assigned to 4 groups: 26 animals (including 5 contact birds introduced 24 hours post-infection) were assigned to group H3 and were infected with 10^3^ EID_50_ H5N8_HP_. 26 animals (including 5 contact birds) were assigned to group H4 and were infected with 10^4^ EID_50_ H5N8_HP_. 21 animals were assigned to group L7 and were infected with 10^7^ EID_50_ H5N8_LP_. 26 animals (including 5 contact birds) were assigned to group L7H3 and were simultaneously infected with a mixture of 10^7^ EID_50_ H5N8_LP_ and 10^3^ EID_50_ H5N8_HP_. 26 animals (including 5 contact birds) were assigned to group L7H4 and were simultaneously infected with a mixture of 10^7^ EID_50_ H5N8_LP_ and 10^4^ EID_50_ H5N8_HP_. 5 animals were assigned to the non-infected control group (NI).

Ducks co-infection experiment: A total of 64 ducks were randomly assigned to 4 groups: 21 animals (including 3 contact birds) were assigned to group H4 and were infected with 10^4^ EID_50_ H5N8_HP_. 16 animals were assigned to group L7 and were infected with 10^7^ EID_50_ H5N8_LP_. 21 animals (including 3 contact birds) were assigned to group L7H4 and were simultaneously infected with a mixture of 10^7^ EID_50_ H5N8_LP_ and 10^4^ EID_50_ H5N8_HP_. 5 animals were assigned to the non-infected control group (NI).

### *In vivo* experimental infections

Serum was collected from all animals pre-infection to ensure that they were serologically negative to influenza virus by using a commercial influenza A NP antibody competition enzyme-linked immunosorbent assay kit (ID Screen; ID-Vet, Montpellier, France) according to the manufacturer’s instructions. When they were 4-week-old, animals were infected through the choanal route using an inoculum volume of 100µL. Non-infected groups received the equivalent volume of allantoic fluid collected from non-infected SPF embryonated chicken eggs. Contact birds were introduced in the poultry isolators 1 day post-infection. Clinical signs were recorded over 8 to 9 days. Oropharyngeal and cloacal swabs were performed on all animals daily. 5 animals from each group were humanely euthanized and necropsied at 1 and 3 dpi. Moribund animals reaching humane termination criteria (as dyspnea, convulsions, severe lethargy) were humanely killed and also necropsied. For each necropsied animal, brain and lungs were collected and stored frozen in TRIzol reagent (Invitrogen, Carlsbad, CA) or stored in 10% neutral formalin.

### H5N8_LP_ and H5N8_HP_ quantification

Cloacal and oropharyngeal swabs from *in vivo* experiments were briefly vortexed in 500µL of sterile PBS and viral RNA was extracted from 200µL using a QiaCube automated platform according to the manufacturer’s instructions (Cador Pathogen QIAcube HT kit; Qiagen, Toronto, Canada). Allantoic fluids from *in ovo* experiments were collected and viral RNA was extracted from 140µL according to the manufacturer’s instructions (QIAamp viral RNA; Qiagen, Toronto, Canada). cDNA was synthesized by reverse transcription of 5µL of RNA, using a HA-specific primer (5’-GTCCTTGCGACTG-3’) and a RevertAid first-strand cDNA synthesis kit (Invitrogen, Thermo Fisher Scientific) according to the manufacturer’s instructions. Influenza nucleic acid load was determined by qPCR using primers targeting either the polybasic or the monobasic HA cleavage site. qPCR was performed in 384-well plates in a final volume of 10µL using a Bravo automated liquid handling platform (Agilent Technologies, Palo Alto, CA) and a ViaaA 7 real-time PCR system (Applied Biosystems, Foster City, CA) at the GeT-TRiX platform (GénoToul, Génopole, Toulouse, France). Mixes were prepared according to the manufacturer’s instructions (iTaq SYBR green PCR, BioRad) with 2µL of cDNA and a final concentration of 0.5µM of each primer. Absolute quantification was performed using a standard curve based on 10-fold serial dilutions of plasmid containing either H5N8_LP_ or H5N8_HP_ HA gene.

### RNA extraction from tissue samples and cDNA synthesis

For each organ, 30mg portions of tissue were placed in tubes with beads (Precellys lysis kit; Stretton Scientific, Ltd., Stretton, United Kingdom) filled with 1mL of TRIzol reagent (Invitrogen, Carlsbad, CA) and mixed for 5s at 6,000rpm three times in a bead beater (Precellys 24; Bertin Technologies, Montigny-le-Bretonneux, France). After TRIzol extraction, the aqueous phase was transferred to a RNA extraction column and processed according to the manufacturer’s instructions (NucleoSpin RNA; Macherey-Nagel GmbH & Co, Germany). cDNA was synthesized by reverse transcription of 500ng of total RNA using either both oligo(dT)18 (0.25µg) and random hexamer (0.1µg) primers or HA-specific primer and a RevertAid first-strand cDNA synthesis kit (Invitrogen, Thermo Fisher Scientific) according to the manufacturer’s instructions.

### Quantitative PCR from tissue samples

Quantitative PCR for the analysis of host genes expression was performed in 384-well plates in a final volume of 10µL using a Bravo automated liquid handling platform (Agilent Technologies, Palo Alto, CA) and a ViiA 7 real-time PCR system (Applied Biosystems, Foster City, CA) at the GeT-TRiX platform (GenoToul, Genopole, Toulouse, France). Mixes were prepared according to the manufacturer’s instructions (iTaq SYBR green PCR, BioRad) with 2µl of 1:20 diluted cDNA and a final 0.5µM concentration of each primer (Table 1). Quantification of influenza virus nucleic acid load in tissues was performed in 96-well plates with a 10µL final volume according to the manufacturer’s instructions (iTaq SYBR green PCR, BioRad), along with 2µL of cDNA and a final 0.5µM concentration of H5N8_HP_ or H5N8_LP_ specific primers. qPCR was performed on a LightCycler 96 (Roche). Relative quantification was carried out by using the 2^-ΔCt^ method. Chickens: RNA levels were normalized with the geometric mean of GAPDH/HPRT1 mRNA levels. Ducks: mRNA levels were normalized with the geometrics means of RPL4/RPL30 mRNA levels in the lungs and RPL4/GAPDH in the brain. Oropharyngeal Mx mRNA expression was performed in 96-well plates with a final volume of 10µL according to the manufacturer’s instructions (iTaq SYBR green one-step, BioRad), with 2µL RNA and a final 0.3µM concentration of each primer. Mx mRNA levels were normalized with GAPDH mRNA levels in chickens and ducks samples using the 2^-ΔCt^ method.

**Table 1:**
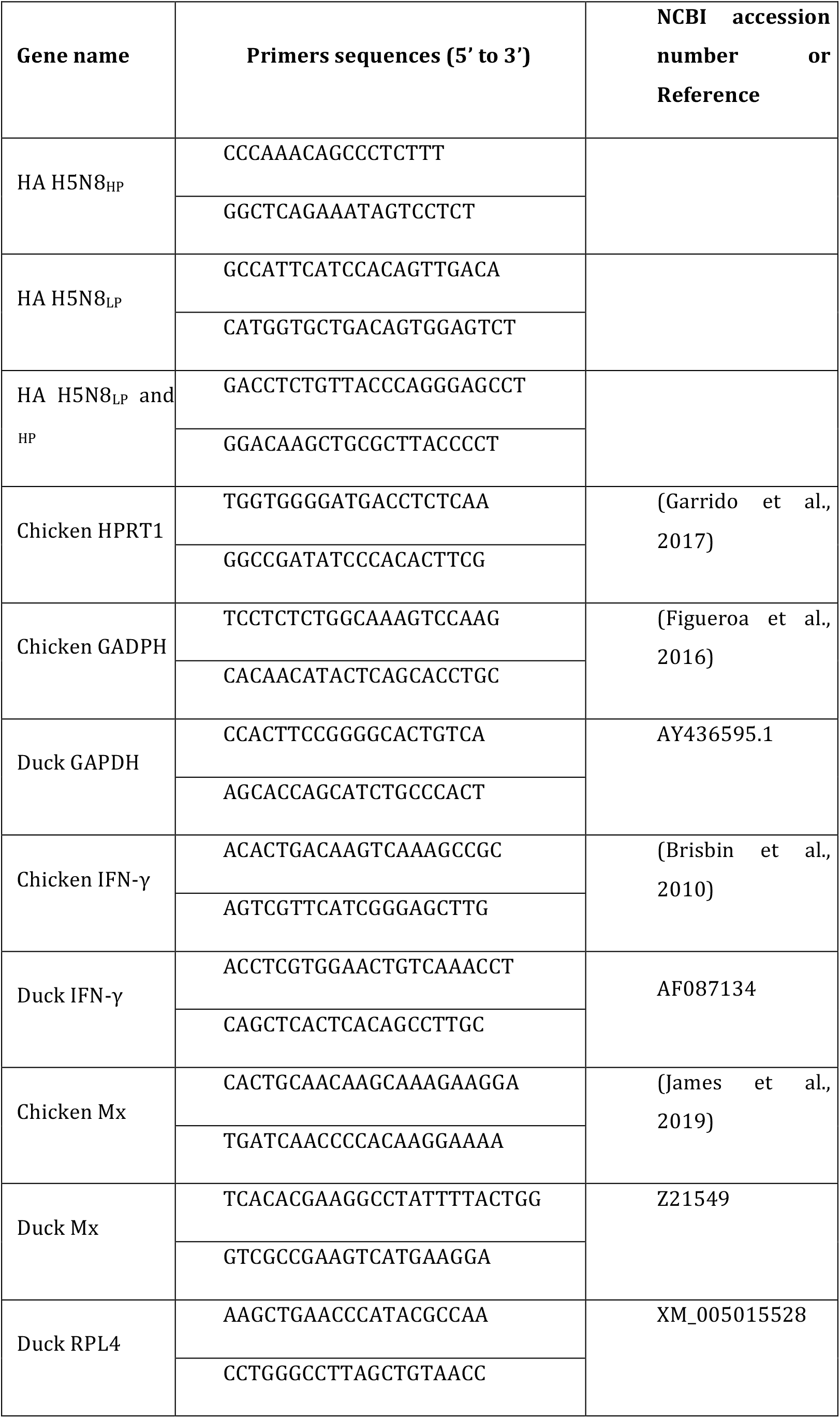
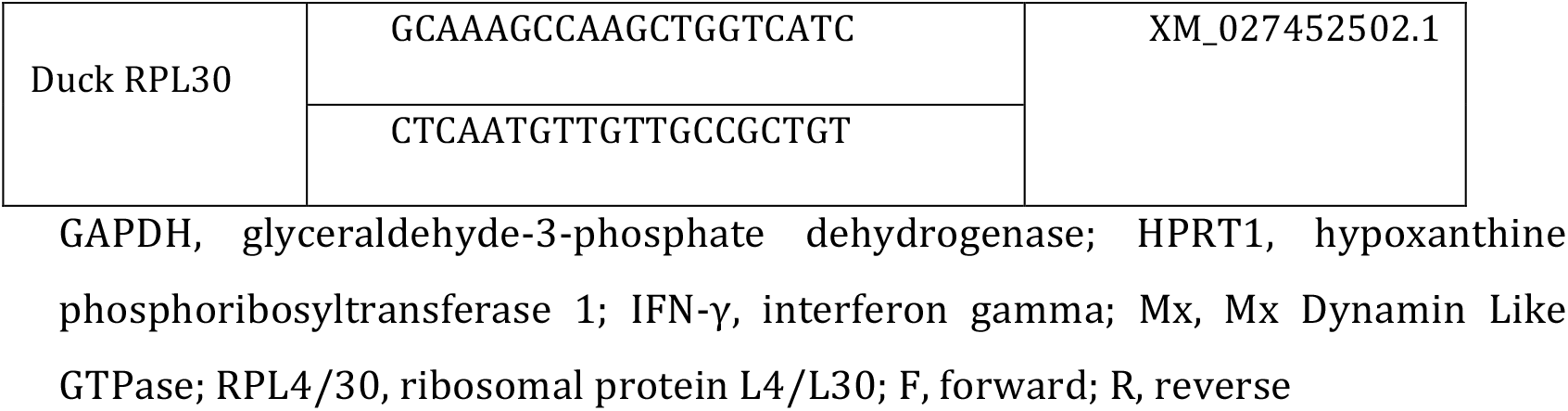
Primers used for qPCR

### Immunohistochemistry

Tissue samples of the lungs and brain were taken and stored in 10% neutral formalin. After fixation, tissues were processed in paraffin blocks, sectioned at 4µm, and immunohistochemistry was performed on paraffin-embedded sections with a monoclonal mouse anti-nucleoprotein influenza A virus antibody (Argene 11-030; pronase 0.05% retrieval solution, 10 min at 37°C: antibody dilution 1/50, incubation overnight at 4°C). The immunohistochemical staining was revealed with a biotinylated polyclonal goat anti-mouse immunoglobulin conjugated with horseradish peroxidase (HRP; Dako, LSAB2 system-HRP, K0675) and the diaminobenzidine HRP chromogen (Thermo Scientific, TA-125-HDX). Negative controls comprised sections incubated either without specific primary antibody or with another monoclonal antibody of the same isotype (IgG2).

## ACKNOWLEDGEMENTS

This work was funded by a grant from the Agence Nationale de la Recherche (ANR-16-CE35-0005-01) to Romain Volmer. Pierre Bessière was supported by a Ph.D. scholarship funded by the Region Occitanie (France) and by the Chaire de Biosécurité at the École Nationale Vétérinaire de Toulouse (French Ministry of Agriculture). Maxime Fusade-Boyer and Gabriel Dupré are supported by PhD scholarships funded by the French Ministry of Research and Education.

## AUTHOR CONTRIBUTIONS

Pierre Bessière, Conceptualization, Formal analysis, Investigation, Visualization, Methodology, Writing – original draft; Thomas Figueroa, Conceptualization, Formal analysis, Investigation, Visualization, Methodology, Writing – review and editing; Amelia Coggon, Charlotte Foret-Lucas, Alexandre Houffschmitt, Maxime Fusade-Boyer, Gabriel Dupré, Jean-Luc Guérin, Maxence Delverdier, Investigation, Writing – review and editing; Romain Volmer, Conceptualization, Formal analysis, Investigation, Visualization, Methodology, Funding acquisition, Writing – original draft, Writing – review and editing, Funding acquisition, Project administration.

## ETHICS

This study was carried out in compliance with European animal welfare regulation. The protocols were approved by the Animal Care and Use Committee (Comité d’Ethique en Science et Santés Animales – 115) under protocol 13025-2018012311319709.

## MATERIALS & CORRESPONDANCE

Correspondence and material requests should be addressed to Romain Volmer.

